# A Reinforcement Learning Model with Choice Traces for a Progressive Ratio Schedule

**DOI:** 10.1101/2023.09.26.559283

**Authors:** Keiko Ihara, Yu Shikano, Sae Kato, Sho Yagishita, Kenji F. Tanaka, Norio Takata

**Affiliations:** Division of Brain Sciences, Institute for Advanced Medical Research, Keio University School of Medicine, 35 Shinanomachi, Shinjuku, Tokyo 160-8582, Japan; Department of Biology, Stanford University, 450 Jane Stanford Way, Stanford, CA 94305-2004, USA; Center for Disease Biology and Integrative Medicine, Faculty of Medicine, The University of Tokyo, 7-3-1 Hongo, Bunkyo, Tokyo 113-0033, Japan

**Keywords:** Choice stickiness, Dopamine, Fiber photometry, Methamphetamine, Mouse, Operant conditioning, Reward prediction error, Ventral striatum

## Abstract

The progressive ratio (PR) lever-press task serves as a benchmark for assessing goal-oriented motivation. However, a well-recognized limitation of the PR task is that only a single data point, known as the breakpoint, is obtained from an entire session as a barometer of motivation. Because the breakpoint is defined as the final ratio of responses achieved in a PR session, variations in choice behavior during the PR task cannot be captured. We addressed this limitation by constructing four reinforcement learning models: a Simple Q-learning model, an Asymmetric model with two learning rates, a Perseverance model with choice traces, and a Perseverance model without learning. These models incorporated three behavioral choices: reinforced and non-reinforced lever presses and void magazine nosepokes (MNPs), because we noticed that mice performed frequent MNPs during PR tasks. The best model was the Perseverance model, which predicted a gradual reduction in amplitudes of reward prediction errors (RPEs) upon void MNPs. We confirmed the prediction experimentally with fiber photometry of extracellular dopamine (DA) dynamics in the ventral striatum of mice using a fluorescent protein (genetically encoded GPCR activation-based DA sensor: GRABDA2m). We verified application of the model by acute intraperitoneal injection of low-dose methamphetamine (METH) before a PR task, which increased the frequency of MNPs during the PR session without changing the breakpoint. The Perseverance model captured behavioral modulation as a result of increased initial action values, which are customarily set to zero and disregarded in reinforcement learning analysis. Our findings suggest that the Perseverance model reveals effects of psychoactive drugs on choice behaviors during PR tasks.

## 1. Introduction

A progressive ratio (PR) schedule in reinforcement learning (RL) is a popular task to measure reward strength (Hodos, 1961; Richardson and Roberts, 1996) and behavioral motivation (Tsutsui-Kimura et al., 2017b; Zhou et al., 2022), but its deficiencies have been well recognized for years (Chen et al., 2022; Richardson and Roberts, 1996). The most significant limitation is that a stream of choice behaviors during the PR session, which commonly takes an hour or more, is discarded, and only a single data point, a breakpoint, is provided from an entire session of a PR task (Arnold and Roberts, 1997). In a PR schedule, response requirements to earn a reward escalate after delivery of each reinforcement, e.g., the number of lever presses required to obtain a single reward increases from 1, 2, 4, … along with trials (Richardson and Roberts, 1996). The highest number of lever presses achieved in a PR session is defined as the breakpoint and is used as a barometer of motivation (Chen et al., 2022). Although modulation of breakpoints by psychostimulants has been used to investigate effects of these drugs (Thompson, 1972), variations in choice behavior during the PR task cannot be captured. Therefore, a method to assess the choice behavior may enable exploration of novel effects of psychostimulants.

Methamphetamine (METH) is a psychoactive dopaminergic drug with a wide variety of effects, including motivational and behavioral effects. Low-dose METH may modulate choice behavior during PR tasks, and this cannot be captured by a breakpoint. Indeed, METH induces qualitatively different effects as a function of dose (Grilly and Loveland, 2001). Moderate doses (1.0–2.0 mg/kg) of METH increase PR task breakpoints (Bailey et al., 2015; Thompson, 1972), but low doses (0.3–0.6 mg/kg) METH have not been reported to exert such modulation (Grilly and Loveland, 2001; Shen et al., 2010). Still, low-dose METH has many other psychological and behavioral effects, including enhancing discrimination of reversal learning (Calhoun and Jones, 1974; Kulig and Calhoun, 1972) and induction of behavioral activation (Hall et al., 2008; Miller et al., 2013) (but see Asami and Kuribara, 1989; Jing et al., 2014). Clinical application of low-concentration dopaminergic drugs for severe post-traumatic stress disorder (PTSD) (Mithoefer et al., 2019), and attention deficit hyperactivity disorder (ADHD) (Guo et al., 2023) further underlines the necessity of developing a quantitative method to enable analysis of behavioral effects by low-dose METH.

Reinforcement learning (RL) algorithms are used to construct normative models that generate subsequent choice behavior based on a history of behavioral selections and rewards (Niv, 2009; Niv et al., 2007). RL models make it possible to relate computation and neurophysiological dynamics, such as encoding of reward prediction error (RPE) by the extracellular dopamine (DA) concentration in the stratum (Schultz et al., 1997). We constructed an RL model for a fixed ratio (FR), lever-press task for mice (Shikano et al., 2023). In the FR schedule, response requirements to earn a reward are fixed (Yokel and Wise, 1975). In our study, we used FR5 tasks that required mice to press a lever 5 times for a reward. To model mouse behavior during FR5 tasks, we constructed an RL model that had multiple state values. Each state value corresponded to a lever-press number, e.g., a state value *V*2 represents a state in which mice pressed a lever twice. Multiple state values in the model assume that mice have an internal representation for each lever press number. It is unlikely, however, that mice possess an internal representation for each lever press in the case of a PR schedule, because the number of lever presses for a reward increases, rapidly exceeding 100. This difficulty may be one of reasons that RL models for PR tasks have apparently not been proposed. A situation in which numerous lever presses are required for mice to obtain a single reward during the latter half of PR tasks resembles a sparse reward environment. A recent study proposed that asymmetric learning rates are necessary for an RL model that describes persistent choice behavior of mice in a scarce reward environment, where the probability for obtaining a reward is small (Ohta et al., 2021). In that study, a large learning rate for a positive RPE, i.e., obtaining a reward, and a small learning rate for a negative RPE, i.e., an unexpected omission of a reward, were proposed as a mechanism for exerting a behavior repeatedly without a reward. Another study, however, demonstrated theoretically that persistent lever pressing behavior is described by an RL model with a choice trace rather than asymmetric learning rates (Katahira, 2018, 2015; Sugawara and Katahira, 2021). It is not clear which model, an asymmetric learning rate model or a choice trace model, better describes choice behavior during PR lever press tasks.

In this study, we propose a RL model with choice traces to realize analysis of choice behavior during PR lever press tasks. We combined a PR lever press task in mice, computational modelling of the behavior, and DA measurements in the ventral striatum (VS) of mice. We found that PR lever-press tasks for mice can be described as a three-choice behavior, rather than two, because mice performed numerous magazine nosepokes (MNPs) to check a food reward, in addition to conventional active and inactive lever presses. A Q-learning model with choice traces was the best-fitting model, as it predicted gradual modulation of RPEs during PR tasks. We confirmed the prediction with DA measurements during the PR tasks by mice. We applied the Perseverance model to experiments with low-dose METH, which did not change breakpoints, but increased MNPs during a PR session. The higher frequency of MNPs during PR tasks was described as increases of initial action values. The Perseverance model realizes examination of choice behavior in PR tasks, which helps to describe effects of psychiatric drugs using PR tasks.

## 2. Materials and Methods

### 2.1. Animals

All animal experiments were approved by the Animal Ethics Committee of Keio University, Japan (approval A2022-301). Eleven 3-month-old, male C57BL/6 mice weighing 23–27-g, purchased from SLC (Shizuoka, Japan), were used. Male mice were used because it is reported that estrous cycle affects performance in PR tasks in rodents (Roberts et al., 1989) and that gender differences exist in behavioral effects of METH, including PR schedules (Roth and Carroll, 2004; Schindler et al., 2002). Mice were housed individually and maintained on a 12-h light/12-h dark schedule, with lights off at 8:00 PM. Their body weights were maintained at 85% of their initial body weight under conditions of food restriction with water ad libitum.

### 2.2 Surgery

Mice were anesthetized by intraperitoneal injection of ketamine (100 mg/kg) and xylazine (10 mg/kg) before a stereotaxic surgery for adeno-associated virus (AAV) injection and implantation of an optic fiber that targeted the right VS (**Supplementary Figure 2**). Details for surgical procedures were described in detail previously (Shikano et al., 2023). Briefly, following an incision in the scalp, a craniotomy with a diameter of 1.5 mm was created above the right VS at stereotaxic coordinates 1.1 mm anteroposterior (AP) and 1.9 mm mediolateral (ML) to the bregma. The dura mater was surgically removed. A total volume of 0.5-µL GRABDA2m virus (PHP.eB AAV-hSyn-GRAB-DA2m-W, 1.0 × 10^14^ genome copies/mL) (Sun et al., 2020, 2018) was injected with a pulled glass micropipette into the VS (3.5 to 3.7 mm dorsoventral (DV) relative to the cortical dura surface) according to the atlas of (Franklin and Paxinos, 2008). The injection was driven at a 100 nL/min flow rate by a microinjector (Nanoliter 2020 Injector, World Precision Instruments, Sarasota, FL). The micropipette was left in place for another 5 min to allow for tissue diffusion before being retracted slowly. Following the GRABDA2m virus injection, an optical fiber cannula (CFMC14L05, 400 µm in diameter, 0.39 NA; Thorlabs, Newton, NJ) attached to a ceramic ferrule (CF440-10, Thorlabs) and a ferrule mating sleeve (ADAF1-5, Thorlabs) was inserted into the same side of the VS as the virus injection and cemented in place (3.4 to 3.6 mm DV). Operant conditioning and data collection were started more than 4 days after the surgery to allow the mice to recover.

### 2.3 Behavioral task

Mice were food-restricted and trained to perform a lever-pressing operant conditioning task in FR- and PR-schedules to retrieve a food pellet, as described previously (Shikano et al., 2023; Tsutsui-Kimura et al., 2017b). Behavioral training and tests were performed under constant darkness in an aluminum operant chamber (21.6 × 17.6 × 14.0 cm; Med Associates, Fairfax, VT) housed within a sound-attenuating enclosure in a daytime. The chamber was equipped with two retractable levers (located 2 cm above the floor), and one food magazine between the levers on the floor (**Fig. 1A**). Each trial began with extension of the levers. As for three mice (mouse ID: VLS06, VLS09, VLS10) of the eleven, a 5-s sound cue (80 dB) from a speaker located on the opposite wall preceded the lever extension. Presses on the lever on the left of the food magazine (reinforced side) were counted (active lever press: ALP), and a reward pellet (20 mg each, Dustless Precision Pellets, Bio-serv, Flemington, NJ) was dispensed to the magazine immediately after the required number of presses was made. The levers were retracted at the same time as the reward delivery. In contrast, presses on the other lever on the right side had no programmed consequence (non-reinforced side; inactive lever press: ILP). A refractory period of 0.5 s followed each lever press before the lever was re-extended. In addition to pressing the reinforced and non-reinforced levers, mice occasionally poked into the magazine (magazine nosepoke: MNP) before making the required number of lever presses (Ko and Wanat, 2016; Shikano et al., 2023; Zhou et al., 2022). The timing of MNP was defined as the time point when the distance of the animal’s head to the center of the magazine became less than 2.5 cm. An inter-trial interval (ITI) of 30 s (or 35 s in the presence of the sound cue) followed each food delivery, during which the levers were not presented, and mice consumed the reward. The subsequent trial was automatically initiated after the ITI period ended. TTL signals were generated at the timings of lever extension and lever pressing and digitized by a data acquisition module (cDAQ-9178, National Instruments, Austin, TX). TTL signals were simultaneously recorded at a sampling frequency of 1,000 Hz by a custom-made program (LabVIEW 2016, National Instruments) using voltage input modules (NI-9215, National Instruments). A single session for the operant conditioning task lasted for 60 min until the mice received 100 food rewards, or when the mice stayed away from the active lever for more than 5 min. To track the moment-to-moment position of the mice, an infrared video camera (ELP 2 Megapixel WEB Camera, OV2710, Ailipu Technology Co., Ltd, Shenzhen, China) was attached to the ceiling of the enclosure. Reflective tapes were attached to a custom-made 3D printed optical fiber protector (1.2 × 1.4 cm) on the head of the mice. The tapes were recorded at a sampling rate of 20 Hz. Mouse positions in each frame were computed offline with a custom-made code (MATLAB 2021a, Mathworks). The entire experimental procedure took 26–32 days, consisting of surgery, recovery, training in FR tasks, and test in PR tasks. Behavioral data were summarized as binary data with an action (ALP, ILP, and MNP), and a reward.

**Figure 1.**
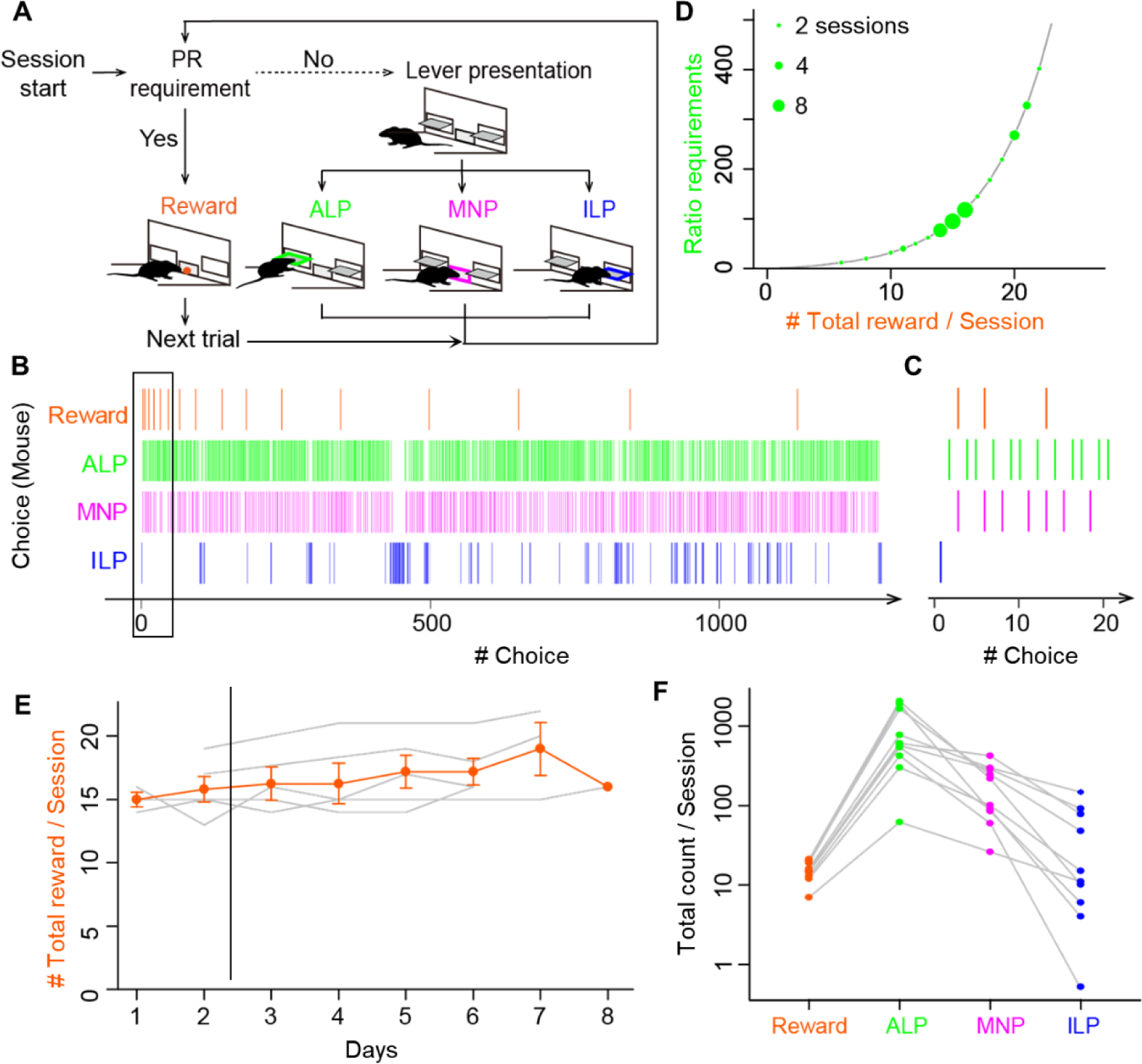
Mice performed void magazine nose pokes as frequently as active and inactive lever presses during progressive ratio lever-press tasks. **A.** Choice behavior by mice during the progressive ratio (PR) lever-press task. A session started with presentation of active and inactive levers (Lever presentation). Mice performed either an active lever press (ALP), a magazine nose poke (MNP), or an inactive lever press (ILP). If mice pressed an active lever a certain number of times (PR requirement), a food reward was delivered at the magazine (Reward), followed by a subsequent trial after an inter-trial interval of 30 s. The number of ALPs required to obtain a reward increased progressively in every trial. **B.** A representative time course of a choice behavior by a mouse in a whole session of a PR task. The session was terminated by a 60 min time-limit, during which the mouse earned 17 food rewards. Intervals of the food rewards (orange) increased due to the exponential increase of the ratio requirement. The mouse checked a magazine without a reward (MNP, magenta) before completing the ratio, in addition to ALP and ILP. **C.** Enlarged view of a time course of a choice behavior (black rectangle in B), showing the first four trials in the session in the PR lever-press task by the mouse. Magazine-checking behavior (magenta) occupied a non-negligible percentage in the choice behavior during the PR task. **D.** The ratio requirement during a PR task. The number of ALP responses required to earn a food reward in a trial increased exponentially in the order: 1, 2, 4, 6, 9, 12, 15, 20, 25, 32, 40, 50, 62, 77, 95, and so on (black line). The y-axis corresponds to breakpoints, which are the final ratios completed in a session. Green circles show breakpoints by 13 mice in 38 sessions. Diameters of circles show the number of sessions by mice. **E.** PR lever-press tasks were stable during Days 3–8. Mice performed the PR task one session per day for 6 to 8 consecutive days. Total reward counts per session (breakpoints) deviated less than 15% after Day 3, data of which were used for the following analysis and modelling. Gray lines show mean reward counts per session of each mouse. **F.** Comparison of total counts of behavioral choices in a session of the PR task. The relative frequency was relatively consistent: ALP > MNP > ILP. Grey lines show data for each mouse. Circles denote average counts of multiple sessions for each mouse (n = 8 mice, 24 sessions).

#### 2.3.1 Fixed- and progressive-ratio tasks

FR sessions were used as a training of mice to associate lever-pressing and a food reward. Mice were required to perform a fixed number of responses (lever presses) to attain a reward: one response was required in an FR1 schedule, and five consecutive responses were required in an FR5 schedule. Mice were trained for at least three sessions (one 60-min session/day) on the FR1 schedule followed by four sessions on the FR5 schedule. FR sessions were finished when the mice accomplished 100 completed trials or spent 60-min for a session. After completing the training using the FR sessions, a lever-press task in a PR schedule started. The operant requirement of each trial increased exponentially following the integer (rounded off) of 5 × exp(*R* × 0.2) − 5, where R is equal to the number of food rewards already earned plus 1 (that is, the next reinforcer), as: 1, 2, 4, 6, 9, 12, 15, 20, 25, 32, 40, 50, 62, 77, 95, and so on (Richardson and Roberts, 1996). The final ratio completed represented a breakpoint (Hodos, 1961).

### 2.4 Computational Models

We constructed four types of RL models (Sutton and Barto, 2018) for a lever-press task in a PR schedule. The model had three behavioral choices based on our experimental findings (**Fig. 1C, F**): a reinforced active lever press (ALP), a non-reinforced inactive lever press (ILP), and a magazine nose poke (MNP). The four models were modifications of a standard Q-learning model (Watkins and Dayan, 1992). (1) The Simple Q-learning model has single state value (hereafter, “SimpleQ”). (2) The Asymmetry model has independent learning rates for positive and negative reward prediction errors (Katahira et al., 2017b; Lefebvre et al., 2017; Ohta et al., 2021). (3) The Perseverance model has a choice auto- correlation to incorporate perseverance in action selection (Katahira, 2018; Lau and Glimcher, 2005). (4) The No-learning perseverance model (“NoLearn”) has a constant learning rate of zero (Katahira et al., 2017a). These models have an action value *Q*^a^_i_, where the subscript *i* is the trial number and the superscript a is for an action a ∈ {ALP, ILP, MNP}. We assigned initial action values *Q*^a^_0_ as free parameters because we assumed that mice would have initial preferences among the choices in the PR task due to pretraining in FR schedules (Katahira et al., 2017a). These models updated an . for a chosen action according to

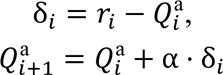

where δ_*i*_ is the RPE, *r*_*i*_is the outcome (reward) at trial *i*, and α is the learning rate, which determines the weight to update action values. The outcome *r*_*i*_ was binarized as 1 for a food reward, and 0 for otherwise. The Asymmetry model had two learning rates α_+_and α_−_ for positive and negative RPEs, respectively. The Perseverance model had additional free parameters *C*^a^_i_ that represent the choice trace for action *α*, which quantifies how frequently action *α* was chosen recently. The choice trace was computed according to the update rule (Akaishi et al., 2014): 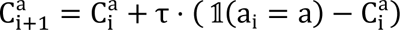, where the indicator function 𝟙(a_i_ = a) assumes a value of 1 if the chosen action *α*_*i*_ at trial *i* is equal to an action a ∈ {ALP, ILP, MNP}. Otherwise, it takes a value of 0. The parameter τ is the decay rate of the choice trace. Initial values for the choice trace *C*^*α*^_0_ were set to zero. The NoLearn model had a constant learning rate α = 0.

The probability of choosing an action *α* by the models at trial *i* was calculated using the softmax function: 

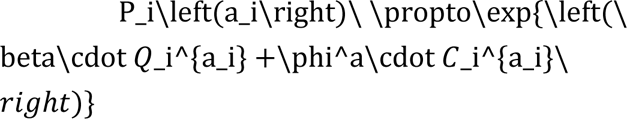

where β is the inverse temperature parameter and ϕ^a^is the choice-trace weight for an action “a” that controls the tendency to repeat (when positive) or avoid (when negative) the action. Only the Perseverance model and the NoLearn model had the parameters, choice trace *C*^ai^_*i*_ and choice-trace weight ϕ^a^

### 2.5 Parameter fitting of these models for behavioral data

Model comparisons were performed based on predictive performance of the models (Palminteri et al., 2017). Maximum log-likelihood estimation was used to fit free parameters of these models to mouse choice behavior during a PR session. The likelihood action value *Q*^a^ for a chosen action according to *L* was calculated with the formula: 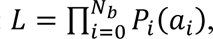 where *N*_*b*_ denotes the last trial number in a PR session, which is equivalent to the ordinal values of a breakpoint. Non-linear optimization was performed to search the most appropriate parameters using the function “optim” in the R programming language. The free parameters *Q*^*α*^and τ had lower and upper bounds from -1 to 1. The Akaike Information Criterion (AIC) was used to compare model fitness to the choice behavior (Daw et al., 2011): AIC = −2 *log*(*L*) + 2 ⋅ *N*_*param*_, where *N*_*param*_is the number of free parameters to fit in the models. Free parameters for SimpleQ, Asymmetry, Perseverance, and NoLearn models were 5, 6, 8, and 7, respectively. The best-fitting parameter values for each model are shown in **Table 1**. The model with the smallest AIC was designated as the best model (Perseverance model).

**Table 1.**
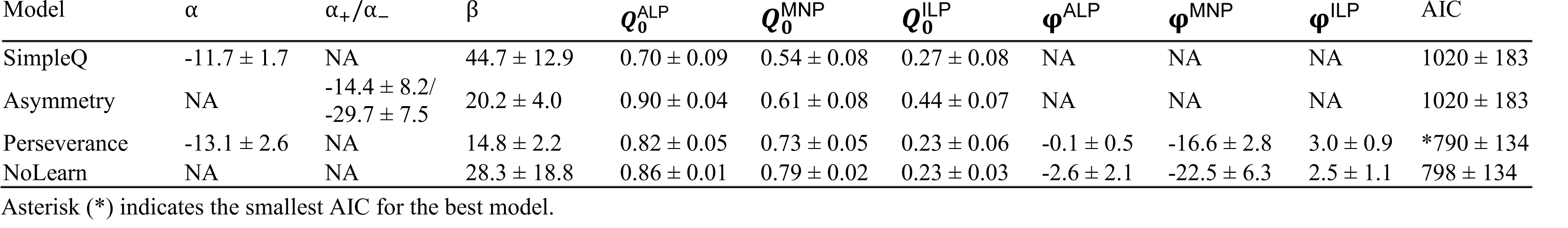
Fitted parameters for reinforcement learning models in lever-press tasks using a progressive ratio schedule in mice.

#### 2.5.1 Model free run

Generative performance of the best model was assessed by a free-run simulation of the three-choice behavior of mice during a PR task using the best model with its best-fitting parameter values (Palminteri et al., 2017; Wilson and Collins, 2019). The last trial number *N*_*b*_ was adopted from a PR session of a mouse K11.

#### 2.5.2 Parameter Recovery and Correlation

Parameter recovery simulation was performed to assess fitting of free parameters of the winning model (Wilson and Collins, 2019). Following the generation of fake choice behavior by the winning model using arbitrary chosen parameter values, we tried to recover the parameters by fitting the best model to the generated data. Association between recovered parameters and true values were checked with Pearson correlation coefficients.

### 2.6 Fiber photometry

Extracellular DA fluctuations were measured using our custom-made fiber photometric system (Natsubori et al., 2017; Shikano et al., 2023). Extracellular DA fluorescence signals were obtained by illuminating cells that expressed GRABDA2m with a 465 nm LED (8.0 ± 0.1 µW at the patch cable tip) and a 405 nm LED (8.0 ± 0.1 µW at the patch cable tip). The 465 nm and 405 nm LED lights were emitted alternately at 20 Hz (turned on for 24 ms and off for 26 ms), with the timing precisely controlled by a programmable pulse generator (Master-8, A.M.P.I., Jerusalem, ISRAEL). Each excitation light was reflected by a dichroic mirror (DM455CFP; Olympus) and coupled into an optical fiber patch cable (400 µm in diameter, 2 m in length, 0.39 NA, M79L01; Thorlabs, Newton, NJ) through a pinhole (400 µm in diameter). The optical fiber patch cable was connected to the optical fiber cannula of the mice. The fluorescence signal was detected by a photomultiplier tube with a GaAsP photocathode (H10722–210; Hamamatsu Photonics, Shizuoka, Japan) at a wavelength of 525 nm. The fluorescence signal, TTL signals that specified the duration of the 465 or 405 nm LED excitations, and TTL signals from behavioral settings were digitized by a data acquisition module (cDAQ-9178, National Instruments, Austin, TX) with a voltage input module (NI-9215, National Instruments). The group of digitized signals was simultaneously recorded at a sampling frequency of 1,000 Hz by a custom-made program (LabVIEW 2016, National Instruments). The fluorescence signal was processed offline, yielding a ratiometric 465/405 signal at a frame rate of 20 Hz, which represented extracellular DA concentration (Shikano et al., 2023). The processed ratiometric signal trace was high-pass filtered at about 0.0167 Hz, corresponding to a wavelength of 1 min to exclude low-frequency fluctuations. We calculated z-scores of the DA signal using the last 20 s (66%) of the ITI period prior to a trial start (**Fig. 3C**) of every trial in a session. Using only the latter part of ITI was important for the calculation of z-scores because DA fluctuation during the first half of ITI may be contaminated by DA fluctuations induced by food consumption. To generate peri-event plots for MNP and Reward, DA fluctuations were binned temporally into blocks of 100 ms.

### 2.7 Histology

After completion of the behavioral task, location of an optical fiber insertion and expression pattern of GRABDA2m protein in the striatum was assessed with a brain slice (**Supplementary Figure 2**). Mice were subjected to the same anesthesia described in the surgery section and were intracardially perfused with 4% paraformaldehyde phosphate buffer solution. Brains were removed and cryoprotected in 20% sucrose overnight, frozen, and cut into 50-μm thick sections on a cryostat (Leica CM3050 S, Leica Biosystems, Wetzlar, Germany). Sections were mounted on silane-coated glass slides (S9226, Matsunami Glass, Osaka, Japan). The GRABDA2m signals received no amplification. Fluorescence images were captured with an all-in-one microscope (BZ-X710, Keyence, Osaka, Japan).

## 3. Results

### 3.1 Void Magazine nose poke is a major behavioral choice in a progressive ratio lever press task

Lever-press tasks for mice have been commonly treated as two-choice tasks with active and inactive lever presses (Ito and Doya, 2015; Tsutsui-Kimura et al., 2017b), but recent studies suggested a void MNP, which is a checking behavior by mice of a food magazine before completing the required number of lever presses, as a third behavioral choice during lever press tasks (Ko and Wanat, 2016; Wanat et al., 2013; Zhou et al., 2022). Indeed, we incorporated an MNP as the third choice in our RL model for a lever press task in an FR5 schedule, demonstrating the relation between an RPE in the model and the frequency of MNP by mice (Shikano et al., 2023). Therefore, we asked if an MNP is also a major behavioral choice in a lever press task in a PR schedule.

Five mice were trained first to associate active lever used for stable performance of PR tasks (**Fig. 1E**). Fluctuation of a breakpoint during day 3 to 8 was smaller than 15% without a significant difference (one-way repeated-measures ANOVA, F[4, 20] = 0.55, p = 0.698). Total counts of MNP (183 ± 41, n = 10 mice, 30 sessions) were comparable to those of other behavioral choices (Reward, 15.2 ± 1.3; ALP, 885 ± 224; ILP, 41 ± 16) (**Fig. 1F**), suggesting that it is important to incorporate MNPs as one of the behavioral choices in a reinforcement learning model for a lever press task in a PR schedule for mice.

### 3.2 Reinforcement learning model with a perseverance factor best replicated a choice behavior during a PR session

We constructed four RL models with three behavioral choices (ALP, ILP, and MNP) to assess choice behavior during a PR lever press task. The agent of the RL models followed the steps in **Fig. 2A**: An agent checked first whether a PR requirement was fulfilled, or a sufficient number of ALPs was performed. If the PR requirement was fulfilled, an agent received a reward and the trial number was incremented, followed by updates of a state value *Q*^MNP^ and a choice trace *C*^MNP^_*i*_ if present. pressing and a food pellet reward with lever-press tasks in FR1 and FR5 schedules. After establishing the association, mice performed a lever-press task in a PR schedule once per day for 6–8 days (**Fig. 1A**). We observed that mice frequently chose an MNP in addition to ALP, and ILP during the PR session (**Fig. 1B**). Mice received a reward pellet from the magazine when a sufficient number of ALPs for a trial were achieved (Vertical orange lines for Reward in **Fig. 1B**). MNPs occurred intermittently, rather than continuously, with other choice behaviors interspersed (**Fig. 1C**). The maximum number of ALPs for one reward in a session, which is defined as a breakpoint, was from 20 to 402, resulting in 8 to 22 pellet rewards in a single session (**Fig. 1D**). In the modeling analysis in the following sections, behavioral data for PR tasks from day 3 to 8 were When the requirement was not fulfilled, an agent chose a behavior among ALP, MNP, and ILP, followed by updates of the state value *Q*_*i*_and the choice trace *C*_*i*_for a chosen behavior reflecting its outcome (no reward). An agent repeated these steps for actual times in a PR session by mice.

**Figure 2.**
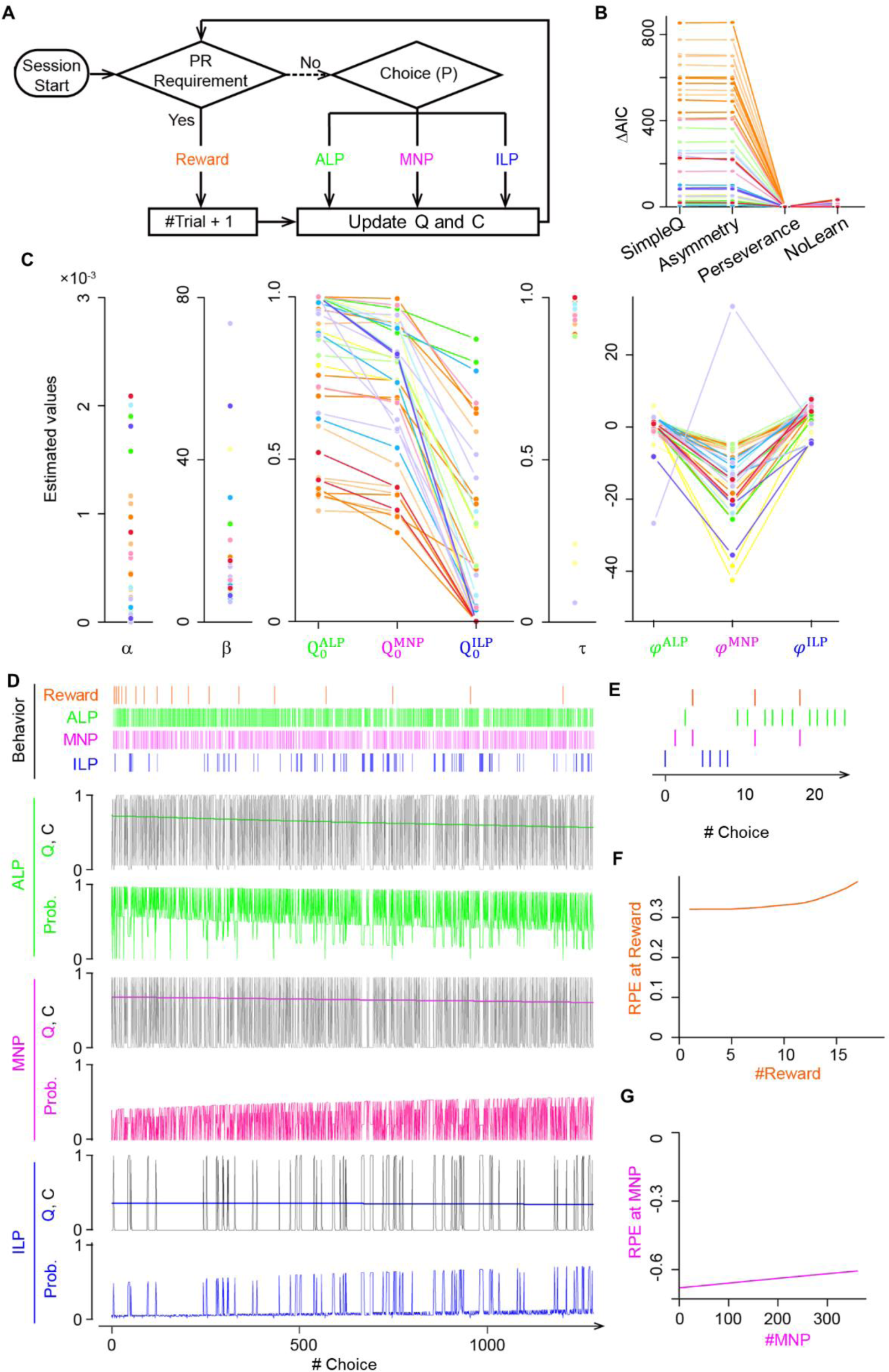
A Q-learning model with choice traces predicted modulation of reward prediction errors during the progressive ratio lever-press task. **A.** PR lever-press task for reinforcement learning models. An agent of the Q-learning models chooses an ALP, a MNP, or an ILP, followed by updates of action values *Q* and/or a choice trace *C* until the PR requirement is met. **B.** Comparison of goodness-of-fit of four Q-learning models for the PR lever-press task: 1) the SimpleQ model with a learning rate *α*, 2) the Asymmetry model with two learning rates, *α*+ and *α*−, for positive and negative RPEs, respectively, 3) the Perseverance model with a learning rate *α* and choice-trace weights *φ*^ALP^, *φ*^MNP^, and *φ*^ILP^ for choice traces *C*^ALP^, *C*^MNP^, and *C*^ILP^, respectively, for modelling the effects of choice hysteresis, and 4) the NoLearn model, which is the perseverance model, with a constant learning rate *α* = 0. AIC was calculated for the four Q-learning models fitted to mouse behavioral data. Lines of the same hue represent data from the same mice. The Y-axis represents differences of AIC from the Perseverance model. The AIC value of the Perseverance model was significantly smaller than that of other models, indicating the Perseverance model as the best model (paired t-test comparing AIC values of the Perseverance model and comparable models: p = 0.003, 0.003, and 0.002 for SimpleQ, Asymmetry, and NoLearn models, respectively). Lines connect the same session of a mouse (38 sessions from 13 mice). **C.** Values of free parameters in the Perseverance model, fitted to mouse behavior. The initial action value for MNP, *φ*^MNP^, was comparable to that for ALP, leading to frequent MNPs by mice during PR sessions. Choice trace weights for MNP, *φ*^MNP^, were significantly smaller than for ALP and ILP, suggesting a lower tendency for consecutive MNP. **D.** Free-run simulation of the Perseverance model. A representative choice pattern (top row) resembles that of actual mice (Fig. 1B). Subsequent rows show timeseries of action values *Q*, choice traces *C*, and choice probabilities for each action ALP (green), MNP (magenta), and ILP (blue) during the simulation of PR sessions. Action values *Q* for ALP (green) and MNPs (magenta) were comparable, reflecting association of lever pressing and a reward delivery. Choice trace *C* modulates the probability (Prob.) of choosing an action by increasing the input values to the softmax function. Probabilities for actions are complementary and sum to 1. **E.** Expanded view of the free-run simulation for the first 20 choices by the Perseverance model. **F, G.** The Perseverance model predicted increasing (**F**) and decreasing (**G**) amplitudes in RPEs upon a reward delivery (Reward, orange) and a magazine nose poke (MNP, magenta), respectively, over the course of the PR session.

The four RL models were 1) SimpleQ, 2) Asymmetry, 3) Perseverance, and 4) NoLearn models (**Fig. 2B**). The SimpleQ model is most commonly used in model-based analysis of choice behavior (Katahira et al., 2017b). The Asymmetry model had distinct learning rates for positive and negative RPEs, respectively. The reason to investigate the Asymmetry model is that repeated behavioral choices required for an ALP in order to obtain a reward in the current PR tasks may resemble scarce environments (Ohta et al., 2021). A recent study demonstrated that rodents in scarce-reward environments had uneven learning rates with a ratio *α*_+_/*α*_−_of about 10, indicating that an agent updates values 10 times more with a positive RPE, i.e., obtaining a reward, than a negative RPE, i.e., no reward (Ohta et al., 2021). Fitting the Asymmetry model to actual mouse behavior during PR tasks resulted in a learning-rate ratio *α*_+_/*α*_−_ of 1.6 ± 1.2 (fitted to behavioral data from 13 mice with 38 sessions; *α*_+_, -14.4 ± 8.2; *α*_−_, -29.7 ± 7.5), which was significantly smaller than the ratio for scarce- reward environments, implying that mice did not regard PR tasks as scarce environments. AIC values of the Asymmetry model were comparable to those of the SimpleQ model, suggesting that introduction of asymmetric learning rates did not increase fitting of the choice behavior of PR lever press tasks (**Fig. 2B**). The third model, the Perseverance model, incorporated a choice trace to represent perseverance small learning rate implied that a Perseverance model with a constant learning rate *α* = 0 (NoLearn model) might be enough for PR tasks. Therefore, we compared AICs for Perseverance and NoLearn models (**Fig. 2B**), obtaining a significantly smaller AIC for the Perseverance model (**Fig. 2B**. Difference of AICs between NoLearn and Perseverance models, 8.4 ± 2.2, p = 0.002). Thus, a small, positive learning rate was necessary to describe PR tasks. In conclusion, the Perseverance model achieved the best predictive performance in the PR lever press tasks for mice (**Fig. 2B**).

Fitted parameters of the Perseverance model to behavioral data of mice are shown in **Fig. 2C** and summarized in **Table 1** with other models, demonstrating a consistent tendency in parameter fitting (**Fig. 2C**. n = 13 mice; learning rate α (6.8 ± 2.1) × 10^-4^; inverse temperature β 14.3 ± 2.3; initial state values *Q*^ALP^ 0.91 ± 0.04, *Q*^MNP^ 0.77 ± 0.04, in action selection (Akaishi et al., 2014; Lau and Glimcher, 2005). The Perseverance model was investigated because a simulation study demonstrated that a model without a choice trace could wrongly assign asymmetric learning rates for perseverance behavior (Katahira, 2018; Sugawara and Katahira, 2021) and because we observed a persistent behavior of mice during PR tasks (**Fig. 1B**). The AIC of the Perseverance model was significantly lower than that of the SimpleQ and Asymmetry models (**Fig. 2B**), implying that repeated ALP in PR tasks are better described as persistence rather than asymmetric learning (difference of AIC values of a model from that of the Perseverance model. P-values obtained by a paired t- test: SimpleQ, 1016 ± 183, p = 0.003; Asymmetry, 1015 ± 183, 0.003; n = 13 mice, n = 38 sessions) (Katahira, 2015; Lau and Glimcher, 2005; Schönberg et al., 2007). The Perseverance model showed stable action values Q during PR sessions (**Fig. 2D**, Q values for ALP, MNP, and ILP), reflecting its small learning rate (**Fig. 2C** *α*). This *Q*^ILP^ 0.69 ± 0.04; decay rate of the choice trace weight τ 0.69 ± 0.04; choice trace weight φ^ALP^ -0.39 ± 0.80, φ^MNP^ -13.9 ± 2.0, φ^ILP^ 3.1 ± 0.6).

Specifically, the initial state value for MNP was significantly larger than that for ILP, further supporting the notion that MNP constitutes a major behavioral choice during lever-press PR sessions for mice (n = 13 mice, t-test, *p* = 2.6 × 10^-6^). Choice trace weights *φ* for MNP were negative (except for a mouse indicated in a grey line in **Fig. 2C**) and significantly smaller than that for ALP or ILP, implying that mice had a tendency to avoid consecutive MNPs.

Generative performance of the Perseverance model was checked by performing a free run of the model with the best-fitting parameter values (Palminteri et al., 2017). Timeseries of simulated choice behavior of the Perseverance model (**Fig. 2D**) were similar overall to actual mouse behavior during PR tasks (**Fig. 1B**). The Perseverance model succeeded in replicating characteristic mouse choice behavior (**Fig. 1D**): repetitive ALPs and continual MNPs with intervals (**Fig. 2E**). We found gradual increases and decreases of RPEs upon reward delivery (**Fig. 2F**) and upon MNPs without reward deliveries (**Fig. 2G**), reflecting gradual decreases of action values for MNPs during PR sessions (*Q*^MNP^ in **Fig. 2D**). These results predict corresponding DA dynamics in the striatum of actual mice because of the proposed relation between RPEs and DA dynamics (Schultz et al., 1997).

We checked the validity of parameter fitting of the Perseverance model with parameter recovery experiments (**Supplementary Figure 1**) (Wilson and Collins, 2019). We used Pearson’s correlation coefficients to confirm that it was able to recover pre-set parameter values by fitting the Perseverance model to the choice behavior sequence obtained by free running the Perseverance model with pre-set parameter values. Correlation coefficients were comparable to previous reports (**Supplementary Figure 1A**: 220 simulations; Correlation coefficients for parameters: α, 0.502; β, 0.383; τ, 0.777; *Q*^ALP^, 0.450, *Q*^MNP^, 0.433; *Q*^ILP^, 0.583; φ^ALP^, 0.670; φ^MNP^, in the brain (Schultz et al., 1997), we measured extracellular DA in the VS of mice during PR lever press tasks to validate predictions of the Perseverance model.

We injected an AAV virus to express a genetically encoded optical DA sensor—GRABDA2m—in the VS (**Fig. 3A**). The VS is involved in lever pressing operant tasks under FR schedules in terms of intracellular calcium activity (Natsubori et al., 2017; Tsutsui-Kimura et al., 2017b, 2017a; Yoshida et al., 2020) and extracellular DA dynamics (Shikano et al., 2023). Our custom optical fiber system enabled monitoring of extracellular DA level fluctuations in the VS during PR sessions (**Fig. 3B**) (Shikano et al., 2023). Ratiometric calculation of GRABDA2m fluorescence signals excited at 405 nm and 465 nm corresponded to extracellular DA dynamics (**Fig. 3C**). Ratio metric calculation helped to distinguish DA decreases and artifactual fluorescence drops due to fluorescence bleaching or body movements. There were large DA increases upon reward delivery and small DA decreases upon MNPs (**Fig. 3C**), consistent with previous studies (Ko and Wanat, 0.545; φ^ILP^, 0.654), suggesting satisfactory parameter recovery (Daw et al., 2011). We also confirmed that there was no significant correlation among recovered parameters (**Supplementary Figure 1B**), suggesting that free parameters in the Perseverance model were independent. These results support the feasibility of parameter fitting and model construction.

**Figure 3.**
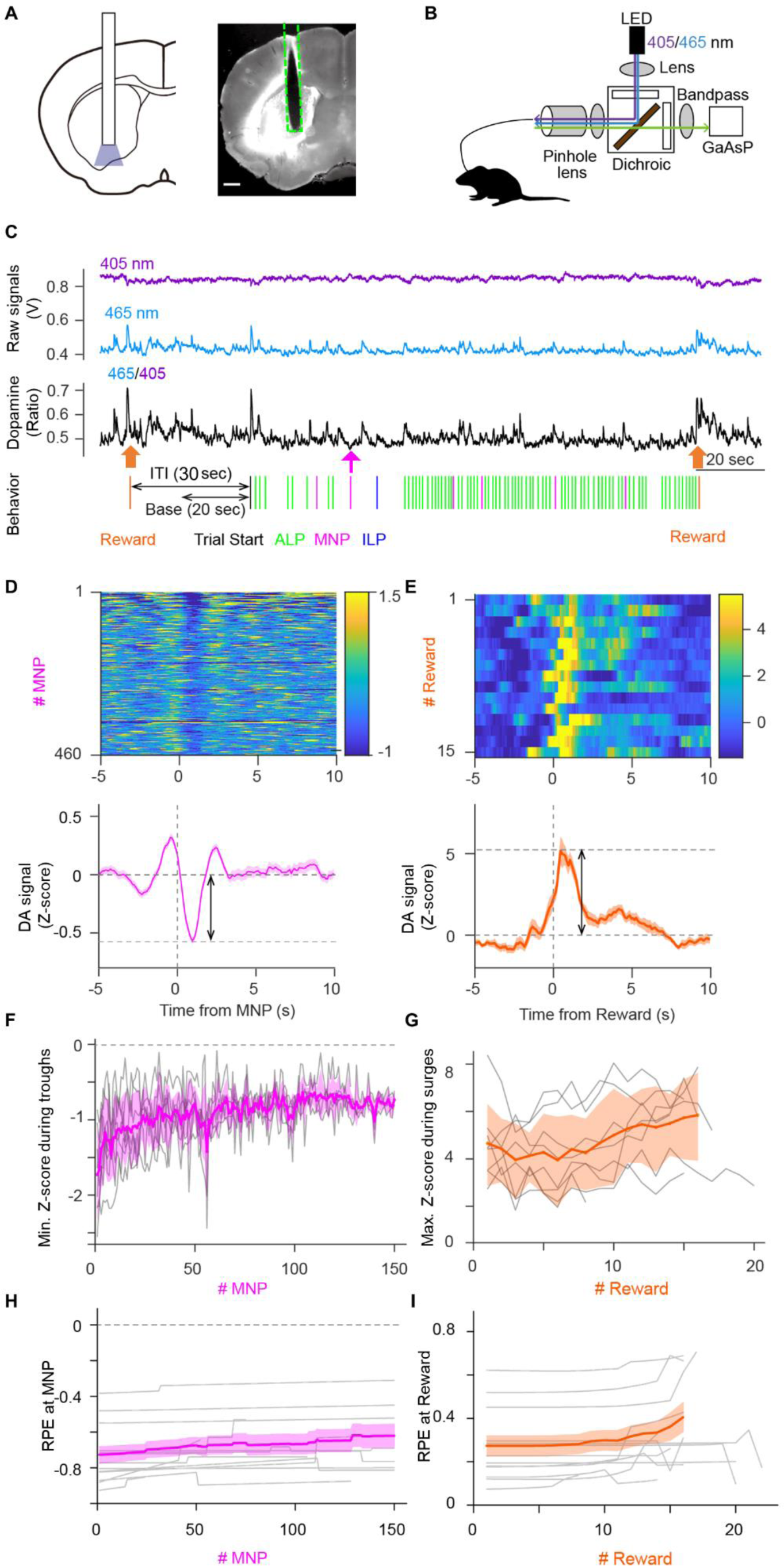
Dopamine dynamics in the ventral striatum during the PR task validated predictions of the Perseverance model. A. Schematic illustration of optical fiber insertion into the VS of mice (Left) and a representative mouse brain coronal section showing the expression pattern of a fluorescent DA sensor, GRABDA2m, (white area) with the optical fiber track (dashed line, Right). Scale bar, 500 µm. B. Fiber photometry system. A single multimode fiber was connected to the optical fiber implanted in the VS. Excitation light was applied continuously with alternating wavelengths of 405 nm (purple) and 465 nm (blue). Fluorescence emissions at 525 nm (green) were detected. C. Representative GRABDA2m signal dynamics during a PR lever press task. Fluorescence intensity fluctuations of GRABDA2m excited by 405-nm (purple) or 465-nm (blue) light were divided to obtain a ratio (465/405, black) as a proxy for extracellular DA concentration. Accompanying choice behavior of the mouse during the PR task is shown at the bottom. Reward delivery (orange arrow) induced a large DA surge. An MNP (magenta arrow) accompanied a small transient DA decrease. The latter part of inter- trial interval (ITI) was used as the Base period to normalize the DA signal. **D, E.** Representative heatmaps showing DA signal fluctuations aligned to MNPs (**D**) or Rewards (**E**) in a session (Top). Averaged DA time courses for heatmaps demonstrate a transient DA decrease at MNP (**D**, bottom), which coincided with the negative RPE upon an MNP in the Perseverance model and a DA surge at Reward (**E**, Bottom), which coincided with the positive RPE upon a reward delivery (**E**, bottom). Double arrows indicate amplitudes of transient DA dynamics. **F, G.** Amplitudes of the transient DA-decrease upon MNPs and the transient DA-surge upon Reward became smaller (**F**) and larger (**G**) during a PR session, replicating the prediction of free-run simulation of the Perseverance model (Fig. 2F, G). Gray lines, DA dynamics of each session of mice. Thick lines in magenta or orange, mean DA dynamics. A transparent area in magenta or orange indicates the standard deviation of DA signal. **H, I.** Time courses of RPEs upon MNPs (**H**) or Rewards (**I**) of the Perseverance model that were fitted to choice-behavior of each session of mice during PR tasks. Slight decreases (**H**) and increases (**I**) of RPEs upon MNP and Reward, respectively, were consistent with the free-run simulation of the Perseverance model (Fig. 2F, G) and with DA dynamics of mice during PR tasks (Fig. 3F, G). Gray lines, RPE dynamics of the Perseverance model fitted to mouse choice-behavior of a session. Thick lines in magenta or orange mean RPEs upon MNPs or Rewards. A transparent area in magenta or orange indicates the standard deviation of RPEs.

### 3.3 Predictions of the Perseverance model on reward prediction errors during the PR task was corroborated by dopamine dynamics in the ventral striatum of mice

Our Perseverance model predicted gradual increases and decreases of RPE-amplitudes upon a reward delivery and a magazine nose poke, respectively, over the course of PR task execution. Because DA is suggested to be a neuronal implementation of RPEs 2016; Shikano et al., 2023). A representative heatmap demonstrated a small, but clear decrease and a large increase in DA fluctuation upon MNP (**Fig. 3D**) and Reward delivery (**Fig. 3E**), respectively. Transient decreases of DA 1-2 s after MNPs are consistent with previous reports (**Fig. 3D** lower panel). Amplitudes of a DA surge upon unconditioned stimulus (Reward delivery) was significantly larger than that of DA decrease upon MNPs, which is also consistent with previous studies (**Fig. 3E** lower panel) (Ko and Wanat, 2016; Shikano et al., 2023).

To examine the model prediction, we plotted time series of DA dip amplitudes upon MNPs (**Fig. 3F**). We performed linear regression to quantify the decreasing trend in DA dip amplitudes, confirming that DA dip amplitudes upon MNP significantly decreased over the PR session (n = 8 mice. Count of MNP, 216 ± 59. Linearly regressed DA dip amplitudes decreased upon an MNP without a reward: slope, (5.4 ± 1.8) × 10^-3^; intercept, -1.1 ± 0.08; t-test on a slope, p = 0.020). We also observed increasing trends of a DA surge upon Reward delivery, although the increasing trend was not significant (**Fig. 3G**; n = 8 mice, session number = 24, Count of a reward supply, 15.1 ± 1.2. Linearly regressed DA surge amplitudes increased upon reward delivery: slope, (5.2 ± 5.5) × 10^-2^; intercept, 4.1 ± 0.5; t-test on a slope, p = 0.374). Fitting of the Perseverance model to behavioral data of mice demonstrated a decrease of DA dip amplitudes upon MNP (**Fig. 3H**) and an increasing trend of DA surge amplitudes upon Reward (**Fig. 3I**). These results support the Perseverance model for describing choice behavior during PR lever press tasks, relating RPEs and DA dynamics in the brain.

### 3.4 The Perseverance model captured effects of low-dose methamphetamine on choice behavior during a PR task as an increase in initial action values

Next, we asked if the Perseverance model can describe modulation of choice behavior during PR tasks by psychiatric drugs. While moderate-dose METH (1.0 mg/kg) injection before lever press PR tasks increased the breakpoint (Bailey et al., 2015; Thompson, 1972), low-dose METH injection has not been reported to change the breakpoint (Asami and Kuribara, 1989; Hall et al., 2008; Jing et al., 2014; Miller et al., 2013). Therefore, behavioral effects of the low-dose drug are not revealed by a breakpoint in PR tasks. To advance computational understanding of modulatory effects of low-dose METH on choice behavior, we applied our Perseverance model to analyze choice behavior of mice during PR tasks.

We first performed behavioral experiments using mice for a lever press PR task. After completing pretraining to associate lever presses with rewards, eight mice each were assigned to Groups A and B for PR tasks for consecutive seven days (**Fig. 4A**). Low-dose METH (0.5 mg/kg i.p.) was injected 10 min prior to a PR session on days 3 and 4 (Group A) or days 6 and 7 (Group B). As control experiments, saline was injected on days 6 and 7 (Group A) or days 4 and 5 (Group B). **Fig. 4B** shows representative choice behavior during a lever-press PR session after Saline (upper panel of **Fig. 4B**) or low-dose METH (lower panel of **Fig. 4B**) injections, demonstrating that low-dose METH increased the frequency of MNPs during a PR session (**Fig. 4B**). We quantified choice behavior during PR tasks after METH or Saline injections, finding that low-dose METH injection significantly increased the number of MNPs in a session (MNPs in **Fig. 4C**, n = 16 mice, t-test, *p* = 0.00012, 117 ± 17 for Saline vs 218 ± 22 for METH) while low-dose METH did not modify the breakpoint (Number of Rewards in a session in **Fig. 4C**. N = 16 mice, session number = 62, t-test, p = 0.12, 16.3 ± 1.0 for Saline, 17.3 ± 0.7 for METH), number of ALPs nor ILPs (**Fig. 4C**. N = 16 mice, t-test; ALP: p = 0.44, count of ALPs, 971 ± 185 for Saline and 1057 ± 150 for METH; ILP: p = 0.16, count of ILPs, 15.3 ± 3.7 for Saline and 23.6 ± 6.1 for METH).

**Figure 4.**
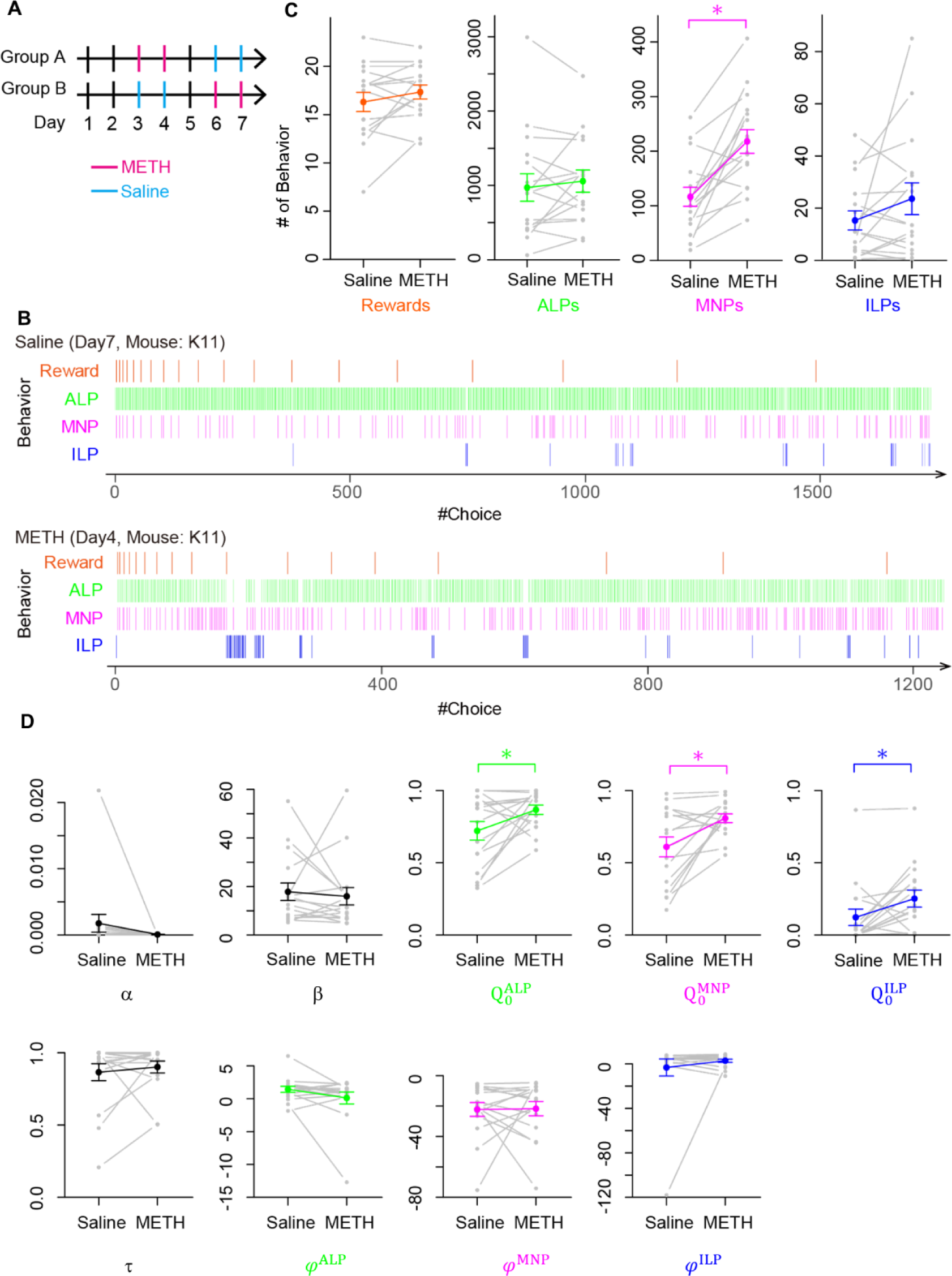
Low-dose METH modified choice behavior during a PR task without changing the breakpoint, which the Perseverance model explained as increases of initial action values. **A.** Schedule of behavioral experiments for a PR lever-press task with methamphetamine (METH, magenta) or saline (blue) injection. Mice in group A received low-dose METH injections (0.5 mg/kg) on days 3 and 4, and saline injections on days 6 and 7. Mice in group B received injections in an opposite manner. On days 1, 2, 5, mice performed a PR task without injection. **B.** Representative time course of choice behavior of mice during a PR lever press task with an injection of Saline (upper) or METH (lower). Slight increase of MNP is visible in a mouse with low-dose METH injection. **C.** Comparison of mouse behavior during a PR session with and without METH injection. Numbers of MNPs increased significantly (n = 16 mice, t-test, p = 0.00012). Other choices including Rewards, which correspond to a breakpoint, were not modified by METH injection (t-test, Rewards, p = 0.12; ALP, p = 0.44; ILP, p = 0.16). **D.** Comparison of free parameters of the Perseverance model that was fitted to choice behavior of mice in a PR session with and without METH injection. Low-dose METH significantly increased initial action values for ALP, MNP, and ILP (n = 16 mice, paired t-test; *Q*^ALP^, p = 0.043; *Q*^MNP^, p = 0.009; *Q*^ILP^, p = 0.031).

We fitted our Perseverance model to the choice behavior of mice during PR tasks after Saline or low-dose METH injection to investigate computational modulation by low-dose METH. We found that low-dose METH increased initial action values for ALP, MNP, and ILP significantly (**Fig. 4D**; n = 16 mice, paired t-test; *Q*^ALP^, p = 0.043, 0.721 ± 0.064 vs 0.866 ± 0.032, *Q*^MNP^, p = 0.009, 0.610 ± 0.068 vs 0.808 ± 0.030, *Q*^ILP^, p = 0.031, 0.123 ± 0.057 vs 0.252 ± 0.059 for Saline vs METH, respectively) while other free parameters were not modulated (n = 16 mice, t-test: α, p = 0.88, -19.2 ± 9.0 vs -20.9 ± 5.0, β, p = 0.68, 17.8 ± 3.6 vs 15.9 ± 3.6; τ, p = 0.52, 0.865 ± 0.059 vs 0.902 ± 0.041; *φ*^ALP^, p = 0.14, 1.43 ± 0.45 vs 0.12 ± 0.90; *φ*^MNP^, p = 0.94, -22.3 ± 4.5 vs -21.8 ± 4.7; *φ*^ILP^, p = 0.42, -3.2 ± 7.7 vs 2.8 ± 1.4). These results imply that low-dose METH increased frequency of MNPs during PR sessions by augmenting initial action values, rather than by learning related free parameters such as α and β.

## 4. Discussion

In this study, we proposed an RL model to analyze choice behavior during a lever press PR task in mice. We demonstrated that choice traces are critical to incorporate perseverance in action selection during PR tasks, rather than asymmetric learning rates. While PR tasks have been widely used to quantify motivation with a breakpoint, this method does not allow assessment of choice behavior during PR sessions because breakpoints are calculated after completing the PR session as the largest number of ALPs achieved for a reward during the session. Our Perseverance model is unique in having a behavioral choice for MNPs in addition to conventional choices for ALP and ILP. Incorporation of MNPs into an RL model was critical in this study because low-dose METH modulated the frequency of MNPs without changing the breakpoint. The Perseverance model predicted a gradual decrease of RPE-amplitudes upon an MNP without a reward delivery during the PR session. We validated the prediction experimentally using fluorescence measurements of extracellular DA in the VS during PR tasks for mice, relating the Perseverance model and neurophysiology. We showed application of the Perseverance model on low-dose METH. The Perseverance model demonstrated that the increase of MNPs during a PR session by METH injection was caused by increased initial action values. The Perseverance model would be a useful tool to investigate effects of psychoactive drugs on choice behavior during lever press PR tasks.

Initial action values were set as free parameters in our model while they are frequently set to zero in RL models (Katahira, 2015). The rationale for setting initial action values to zeros is that agents of RL models choose actions and update action values repeatedly, which decreases the contribution of initial action values to zero asymptotically. In the present study, however, initial action values were indispensable to capture the effect of low-dose METH on a choice behavior during PR tasks. The reason for the significance of initial action values in the present study would be the small learning rate. Due to small learning rates, action values did not change dramatically by value updates upon a choice behavior during a PR session. It is noteworthy that small, but positive learning rates were still necessary for describing choice behavior. The small, but positive learning rate in the Perseverance model allows the model to adapt to a situation in a PR task in which the requirement for lever press counts increases exponentially during a session. Therefore, we presented a unique situation in an RL model for mice, in which modulation of initial action values, rather than learning-related parameters such as a learning rate, resulted in bias in choice behavior (Biele et al., 2011). It is intriguing that not only the initial action value for MNP but also that for ALP and ILP were increased by low-dose METH. These results may coincide with previous reports that acute amphetamine disrupted discrimination of cues with different reward sizes (Werlen et al., 2020). The neurophysiological substrate for increased initial action values is not clear in this study. Increases of tonic DA concentration by METH injection might be related to initial action value modulation because we have shown that moderate- and high-dose METH injection increases extracellular DA concentration in the mouse VS over an hour (Iino et al., 2020). Functional magnetic resonance imaging (fMRI) might be fruitful to reveal brain regions affected by low-dose METH injection (Taheri et al., 2016; Weafer et al., 2019; Yoshida et al., 2016). An fMRI study on human subjects suggested encoding of RPE at the striatum and action values at the medial prefrontal cortex, respectively (Bernacer et al., 2013).

One limitation of the Perseverance model is that it does not describe a breakpoint in PR tasks. RL models including our Perseverance model are suited for describing choice behavior, but commonly do not have a mechanism to stop choosing a behavior. A previous study incorporated a motivation factor in an Actor-Critic RL model for describing thirstiness of the agent in a visual GO/NOGO lick task for mice, succeeded in describing a response rate decrease in the late phase of a session (Berditchevskaia et al., 2016). The motivation factor was an additional positive value to bias an action value for GO choice, which diminishes every time a water reward is received. In that study, a decrease of a GO choice and an increase of a NOGO choice during the late phase of a session was described successfully by the motivation factor. However, in the current PR tasks, mice stop making behavioral choices at a breakpoint. Therefore, simple introduction of the motivation factor into the current Perseverance model cannot identify breakpoints. Mathematical Principles of Reinforcement (MPR) models have been proposed to describe relationships between response requirements and subsequent behavioral pauses and so on, enabling prediction of breakpoints based on regressed parameters (Bradshaw and Killeen, 2012; Killeen et al., 2009). The MPR model is, however, a descriptive model that does not provide a computational basis for each choice behavior in PR tasks. Another caveat is dose-dependent effects of METH. High doses METH (3.0–10.0 mg/kg) induces stereotyped behavior in mice such as focused sniffing, licking, or grooming (Kelley, 2001; Mason and Rushen, 2008; Shen et al., 2010), not performing choice behavior for PR tasks (Hadamitzky et al., 2012; Randrup and Munkvad, 1967). The Perseverance model cannot be applied to such situations. In conclusion, we highlighted the importance of MNPs as a behavioral choice during PR tasks that indicates mouse expectation of a reward. Our Perseverance model analyzes choice behavior during a PR task that expands utility of PR tasks and advances understanding of computational mechanisms of effects by psychoactive drugs that cannot be revealed with a breakpoint.

## Data availability statement

The raw data supporting the conclusions of this article will be made available by the authors, without undue reservation.

## Acknowledgments

We thank T. Fukai and H. Ohta for their advice on the computational modeling.

## Author contributions

KI conceived the experiments based on the advice from KFT and NT. KI, YS and SK conducted the experiments. NT constructed experimental devices and measurement software. KI and NT developed the computational models. KI analyzed data and performed simulation with support from NT. SY prepared the GRAB virus. NT and KI wrote the manuscript. All authors contributed to the article and approved the submitted version.

---

**Supplementary Figure 1.**
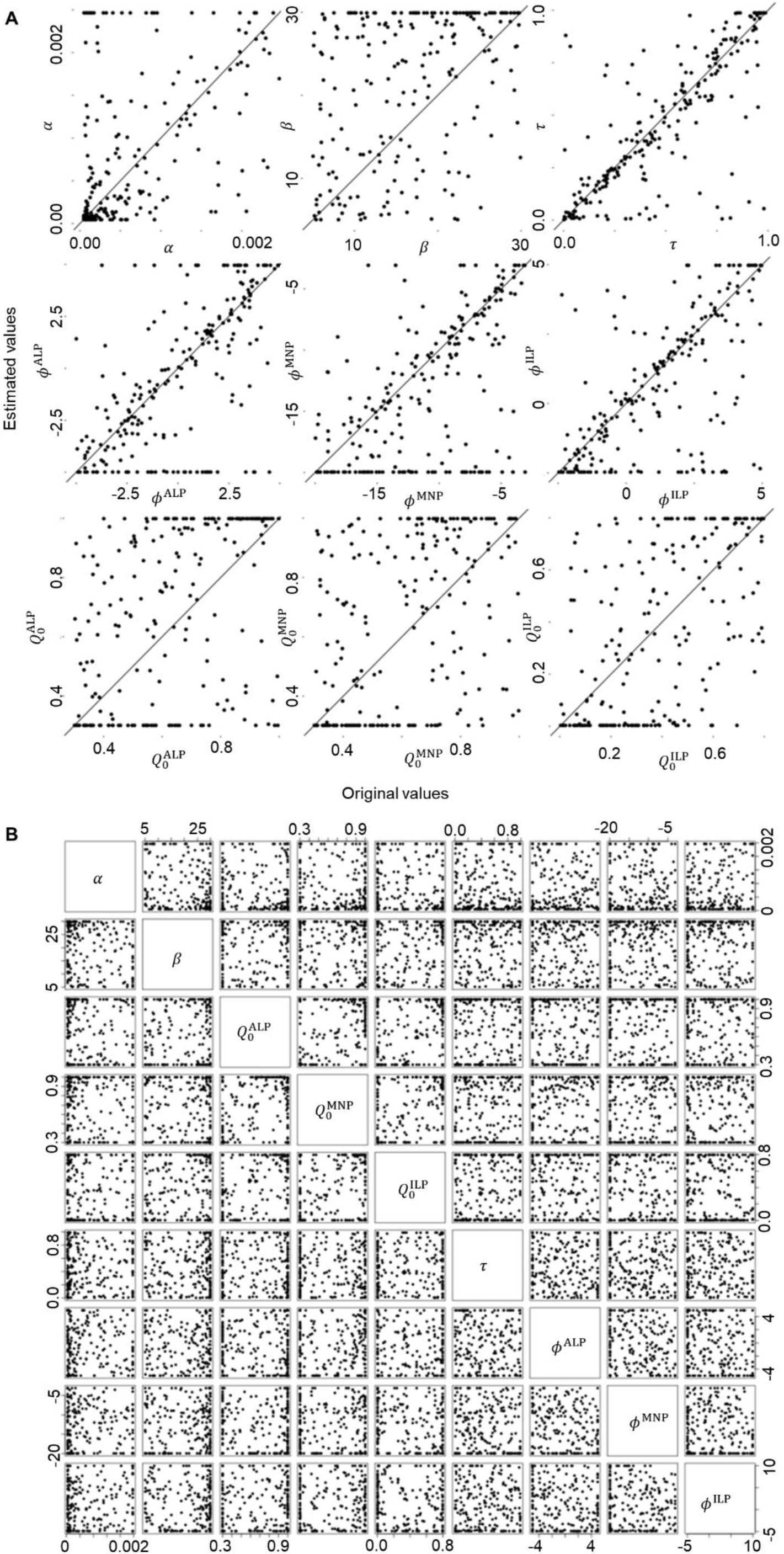
Parameter recovery and correlation of the Perseverance model. **A.** Parameter recovery of the Perseverance model. The Perseverance model was fitted to the choice behavior that was generated using a Perseverance model with arbitrary chosen free parameters for 1,000 sessions. Correlation coefficients between chosen and fitted parameters were comparable to reported values (Daw et al., 2011), justifying our parameter fitting: α, 0.50; β, 0.38; τ, 0.78; *Q*^ALP^, 0.45, *Q*^MNP^, 0.43, *Q*^ILP^, 0.58, *φ*^ALP^, 0.67; *φ*^MNP^, 0.54; *φ*^ILP^, 0.65. **B.** Correlation between estimated parameters in the Perseverance model to assess independence of the parameters. Correlation coefficients were acceptably small (0.0023 to 0.26) implying independence between free parameters.

**Supplementary Figure 2.**
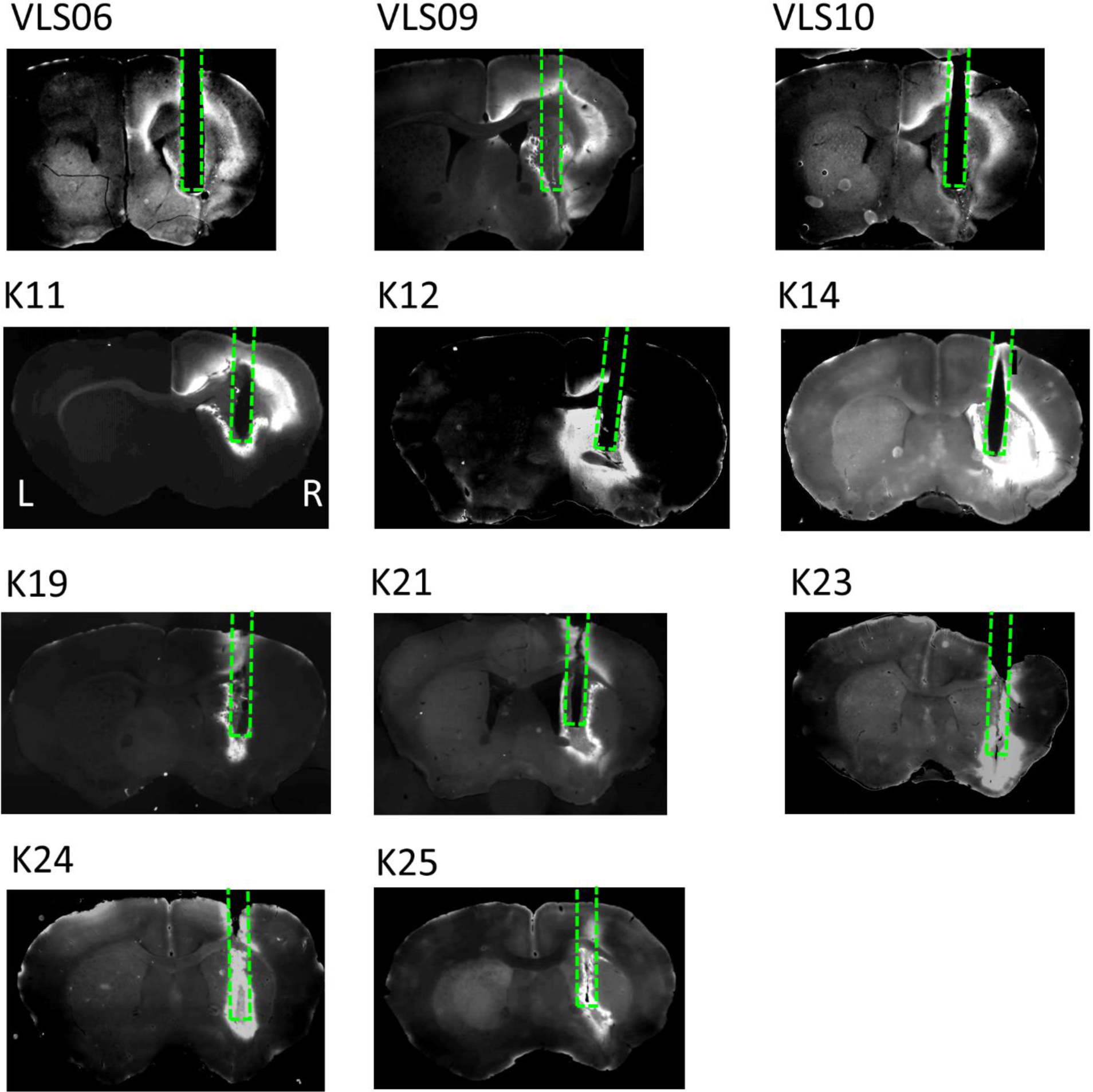
Mouse brain sections showing DA sensor expression and an optic fiber track. Coronal brain sections of mice showing expression pattern of the DA sensor, GRABDA2m, (white area) and the insertion track of the optic fiber targeted at the VS. Animal IDs are indicated at the upper left of each image. The dashed line shows an optical fiber track. Scale bar, 500 µm.

